# CLASP stabilizes microtubule plus ends created by serving to drive cortical array reorientation

**DOI:** 10.1101/196329

**Authors:** Jelmer J. Lindeboom, Masayoshi Nakamura, Marco Saltini, Anneke Hibbel, Ankit Walia, Tijs Ketelaar, Anne Mie C. Emons, John C. Sedbrook, Viktor Kirik, Bela M. Mulder, David W. Ehrhardt

## Abstract

Central to building and reorganizing cytoskeletal arrays is the creation of new polymers. While nucleation has been the major focus of study for new microtubule generation, severing has been proposed as an alternative mechanism to create new polymers, a mechanism recently shown to drive the reorientation of cortical arrays of higher plants in response to blue light perception. As severing produces new plus ends behind the stabilizing GTP-cap, an important and unanswered question is how these are stabilized *in vivo* to promote net microtubule generation. Here we identify the conserved protein CLASP as a potent stabilizer of new plus ends created by katanin severing and find that CLASP is required for rapid cortical array reorientation. In *clasp* mutants both rescue of shrinking plus ends and the regrowth of plus ends immediately after severing are reduced, computational modeling reveals that it is the specific stabilization of severed ends that explains CLASP’s function in promoting microtubule amplification by severing and cortical array reorientation.

## Introduction

Many animal cells, and all higher plant cells, build interphase microtubule arrays with distinct architectures without benefit of a centrosome or other central organizer. In higher plants, interphase microtubules associated with the plasma membrane are organized into highly ordered arrangements oriented transversely to the main axis of cell growth. These cortical arrays are crucial for determining the direction of cell expansion (Baskin, 2001; Wasteneys, 2002; Ehrhardt and Shaw, 2006), in part by guiding the deposition and thus orientation of cellulose as it is synthesized by protein complexes in the plasma membrane (Paredez et al., 2006). New microtubules in these arrays are created by nucleation from γ-tubulin complexes distributed along the sides of existing microtubules (Wasteneys and Williamson, 1989; Chan et al., 2003; Murata et al., 2005; Nakamura et al., 2010) and by a second, recently discovered mechanism – the creation of new microtubule ends by katanin mediated severing at locations where cortical microtubules intersect and crossover each other (Wightman and Turner, 2007; Zhang et al., 2013; Lindeboom et al., 2013b). This severing mechanism for microtubule generation plays an essential role in cortical array reorganization in response to light, a key environmental cue for plant development (Lindeboom et al., 2013b). When seedlings develop in the dark, cells in the embryonic axis, also known as the hypocotyl, undergo rapid axial elongation to push the seedling shoot into the light from the soil. Upon perception of blue light, principally through phototropin photoreceptors, transverse cortical arrays in hypocotyl epidermal cells undergo a remarkable reorientation to a transverse direction, a remodeling that takes place on the scale of approximately 15 minutes. This reorganization occurs by upregulation of katanin mediated severing at crossovers, with a preference for severing of the newer of the two microtubules (Zhang et al., 2013; Lindeboom et al., 2013b). The repeated process of severing and growth of the new plus ends amplifies a new population of microtubules at 90° to the existing array. A functional consequence of this reorientation is redirection of the trajectories of cellulose synthase complexes as they build the cell wall, thus remodeling cell wall structure.

Creation of new microtubules by severing and rescue has also been suggested in animal cells, particularly in neurons, epithelial cells and meiocytes (Roll-Mecak, 2013). A critical question in all these systems is how the new plus ends created by severing are stabilized in order for severing to efficiently build a population of new microtubules. When new microtubules arrays are built by nucleation, the growing ends assembled at nucleation complexes are stabilized by the GTP state of newly added tubulin dimers (Mitchison and Kirschner, 1984) and by proteins that associate with growing ends (Akhmanova and Steinmetz, 2015), some of which are recruited by the GTP-cap (Akhmanova and Steinmetz, 2015). However, new plus ends generated by severing are likely created behind the GTP-cap. When created *in vitro*, these new plus ends shrink immediately (Walker et al., 1989; Tran et al., 1997). By contrast, when severing creates new plus ends *in vivo*, they frequently initiate new growth without observable shrinkage (Lindeboom et al., 2013b). The factors that act *in vivo* to stabilize new plus ends created by severing to permit generation of new microtubules are not known. Here we identify CLASP as a specific rescue factor for new plus ends generated by katanin-mediated severing in the model plant *Arabidopsis*, and demonstrate through quantitative imaging and computational modeling that plus end stabilization by CLASP following severing contributes to an effective reorganization of the microtubule array in response to environmental signals.

## Microtubule reorientation in +TIP mutants

Several proteins that accumulate at and track growing plus ends (+TIPs) have been identified in the model plant *Arabidopsis thaliana* (Bisgrove et al., 2004). We asked if one or more of these +TIPs might act as a specific plus end rescue factor after severing to support creation of a new cortical array. Specifically, we assessed End Binding Protein 1 (EB1) (Mathur et al., 2003; Chan et al., 2003), Cytoplasmic-Linker-Associated Protein (CLASP) (Ambrose et al, 2007; Kirik et al, 2007) and SPIRAL1 (SPR1) (Nakajima et al., 2004; Sedbrook et al., 2004). EB1 and CLASP are both conserved across plants and animals. EB1 is encoded by three genes in *Arabidopsis, EB1a-c*; all three loci are disrupted in the triple mutant *3x-eb1* (Galva et al., 2014). CLASP is represented by a single gene in *Arabidopsis*, where loss of function (*clasp*) results in significant reduction of cell growth anisotropy (Ambrose et al., 2007; Kirik et al., 2007). SPR1 is specific to algae and plants (Furutani et al., 2000). In *Arabidopsis*, SPR1 is part of a six-gene family where loss of SPR1 function itself (*spr1*) causes pronounced right-handed chiral growth of plant organs and loss of cell growth anisotropy. As a first step, we asked if loss of function in any of these candidate proteins were impeded in blue light stimulated cortical array reorganization. Time lapsed imaging of microtubule organization was accomplished by introducing the tubulin marker 35S-YFP-TUA5 into the *spr1, clasp* and *3x-eb1* mutants (Fig. 1A and Video 1), and quantitation of local microtubule orientation over time was performed with the ImageJ plugin LOCO (Lindeboom et al., 2013b) (Fig. 1B and C). We found that both the *3x-eb1* and *clasp* mutants had a higher degree of transverse order than WT before the start of blue light excitation (*P* < 0.05 Mann-Whitney U test compared to WT) (Fig. 1D). Importantly, when the speed with which longitudinal order was built up over time was measured using the ImageJ plugin LOCO (Lindeboom et al., 2013b), the *3x-eb1* and *clasp* mutants were in addition significantly slower than WT (*P* < 0.05 Mann-Whitney U test compared to WT) (Fig. 1E). By contrast, although *spr1* mutants have pronounced morphogenetic phenotypes, no significant effect was measured in either initial array order nor reorientation speed. Thus, both the *3x-eb1* and *clasp* mutants show significant impairment in light-stimulated cortical array reorientation, starting off with higher degrees of transverse order while at the same time showing a slower generation of longitudinal microtubule order.

**Figure 1.**
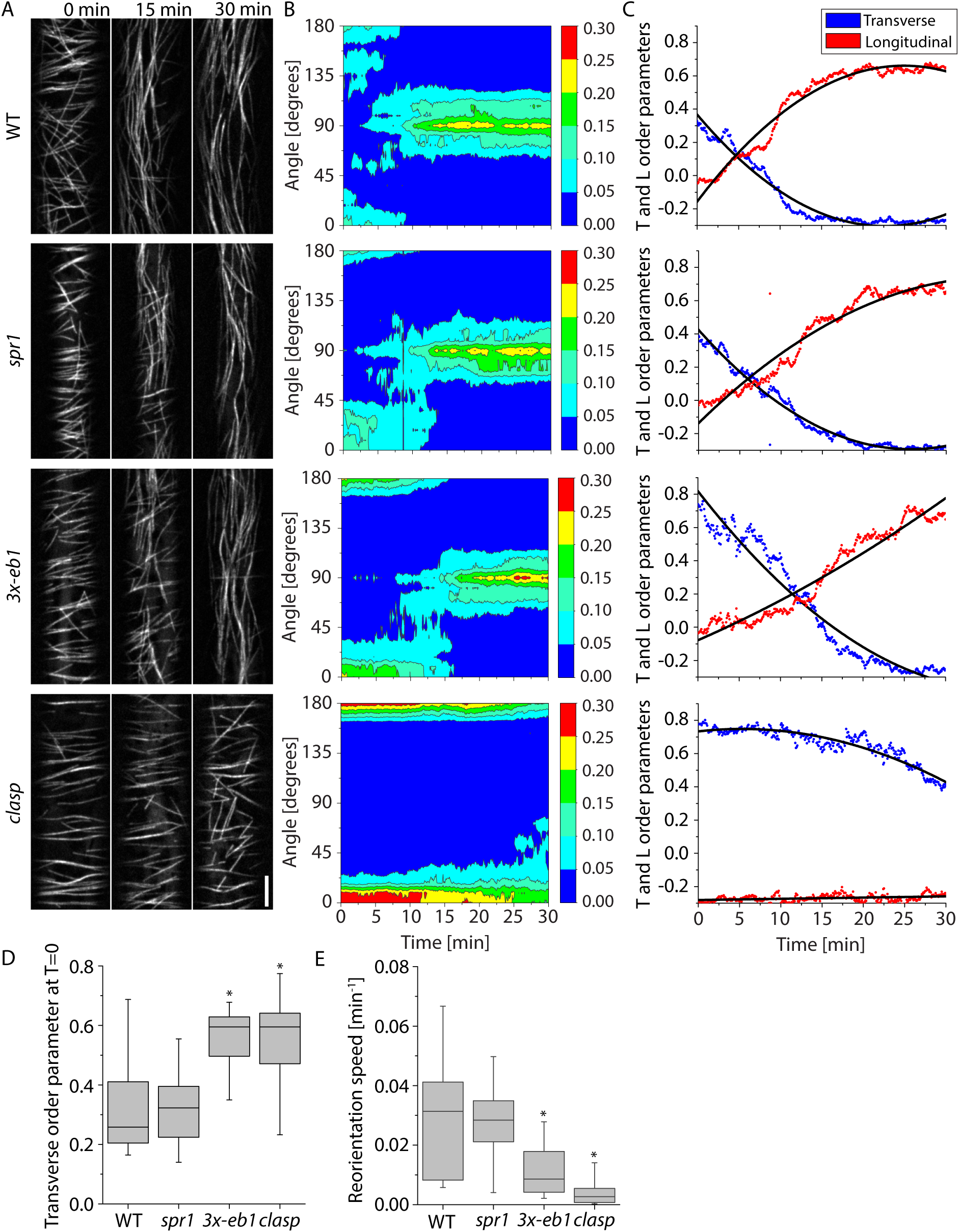
Microtubule reorientation in +TIP mutants. (A) Panels from confocal image time series of dark grown hypocotyl cells expressing YFP-TUA5 in WT, *spr1, 3-eb1* and *clasp* mutant backgrounds at 0, 15 and 30 minutes after induction of reorientation by blue light. See Video 1. Scale bar is 5 μm. (B) Representative contour plots of MT orientation over time corresponding to the movies shown in (A). The color scale represents fraction of microtubules. (C) Longitudinal and Transverse order parameters during MT reorientation for the movies shown in (A). Black lines show quadratic fit. (D) Transverse order parameter at T = 0 for cortical microtubules imaged in WT, *spr1, 3x-eb1*, and *clasp* seedlings. N=8, 9, 9 and 9 cells respectively. A Kruskal-Wallis test showed significant differences among the genotypes (*P* < 0.01). Asterisks represent significant difference from WT by Mann-Whitney U test (*P* < 0.05). (E) Longitudinal reorientation speed (Lindeboom *et al*, 2013b) of imaged microtubule arrays in etiolated hypocotyl cells expressing YFP-TUA5 in WT, *spr1, 3x-eb1*, and *clasp* seedlings. A Kruskal-Wallis test showed significant differences for reorientation speed distributions among the genotypes (*P* < 0.01). N=8, 9, 9 and 9 cells respectively. Asterisks represent significant difference from WT in Mann-Whitney U test (*P* < 0.05). Boxplots show the 25^th^ and 75^th^ percentile as box edges, the line in the box indicates median value and the whiskers show the 2.5^th^ and 97.5^th^ percentile.

The reorientation defects we found for the *3x-eb1* and *clasp* mutants may be due to specific effects on plus end rescue after severing, to changes in basic microtubule dynamics, or possibly severing itself. To assess these possibilities, we first measured individual microtubule dynamics, severing rates at microtubule crossovers, and specific rescue of plus ends after severing at the onset of reorientation. We then explored the consequences of measured changes in microtubules behaviors with modeling and computer simulation studies.

## Microtubule dynamics in +TIP mutants

To measure basic microtubule dynamics, we generated kymographs from the reorientation movies (Fig. 2A) and traced the trajectories of microtubule plus ends (see Methods). In *Arabidopsis* cortical arrays, plus ends are easily distinguished from minus ends by their distinct dynamics, with plus ends showing extended episodes of growth while minus ends show primarily slow shrinkage and pause (Shaw et al., 2003). We observed quantitatively small yet statistically significant differences in growth velocities between WT and all the tested +TIP mutants (*P* < 0.001 Mann-Whitney U test compared to WT for all genotypes) (Fig. 2B, Table S1). *spr1* and *clasp* showed higher growth velocities than WT, while plus ends in *3x-eb1* grew more slowly than WT. We found that *spr1* mutants tended to have slower shrinkage velocities than WT (*P* < 0.05 Mann-Whitney U test), while *3x-eb1* and *clasp* mutants both had higher shrinkage velocities than WT (Fig. 2C, *P* < 0.05 and *P* < 0.01, respectively, Mann-Whitney U test). When making transitions between shrinking to growth we observed that the *clasp* mutants had a significant rescue defect compared to WT (Fig. 2D, *P* < 0.05 rate ratio exact test). No significant differences were measured in catastrophe rates between WT and any of the +TIP mutants (Fig. 2E). In summary, we found small but significant differences in microtubule dynamics between WT and *spr1, 3x-eb1* and *clasp*, the most notable being the rescue defect in *clasp*. The possible significance of these small but significant differences to the amplification of new microtubule array will be explored below in computer modeling studies.

**Figure 2.**
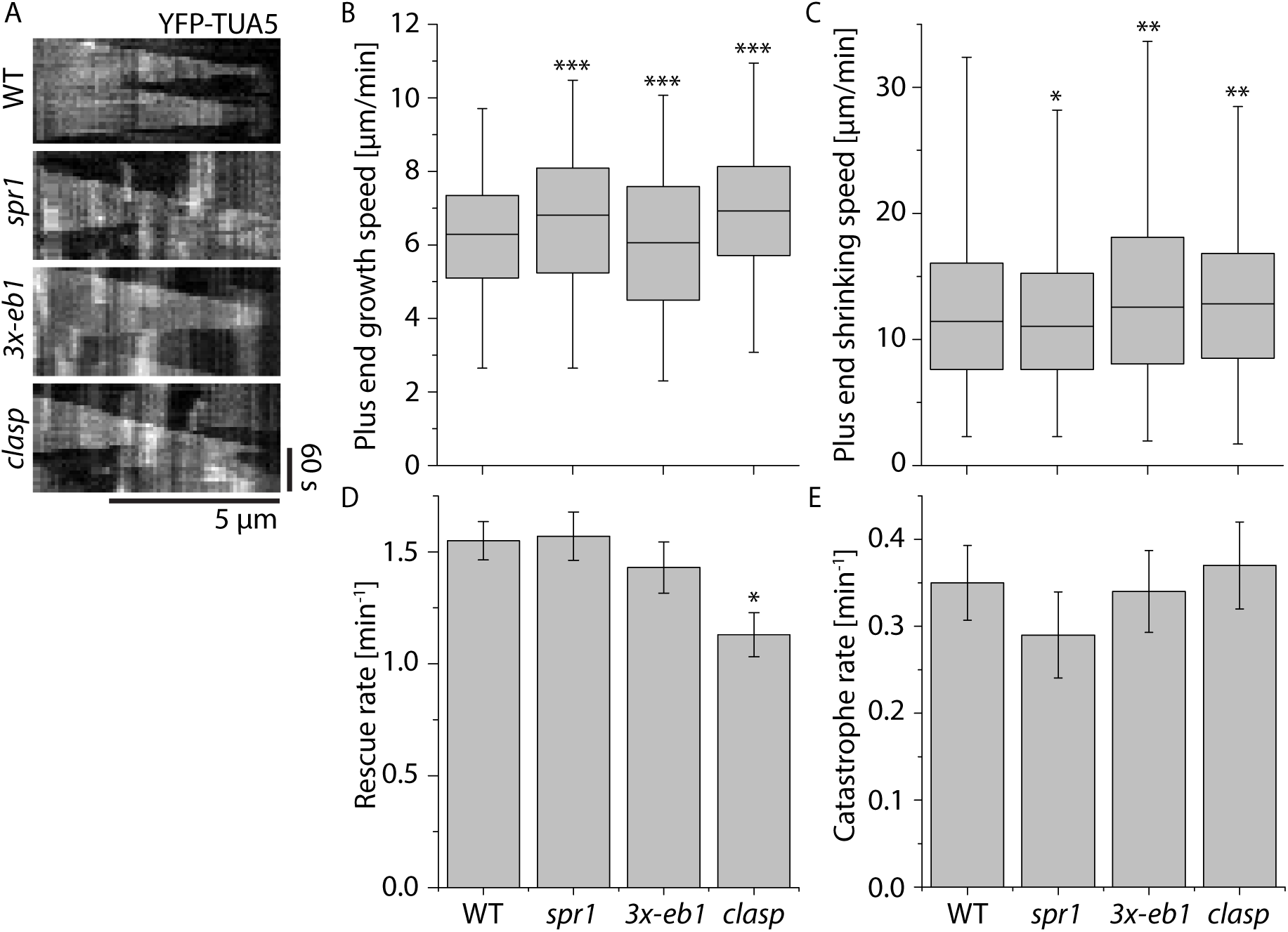
Microtubule plus end dynamics in +TIP mutants. (A) Kymographs of microtubules during early stages of cortical array reorientation induced by blue light in WT, *spr1, 3x-eb1* and *clasp* seedlings expressing YFP-TUA5. (B) Boxplot of microtubule plus end growth speeds and (C) plus end shrinkage speeds in N = 2148, 1405, 1837, and 1516 segments measured in WT, *spr1, 3x-eb1* and *clasp* seedlings respectively. Asterisks indicate a significant difference from WT. (D) Microtubule rescue rates. N = 213, 135, 110, and 116 rescues in WT, *spr1, 3x-eb1* and *clasp* mutant backgrounds respectively. Error bars represent SEM. Asterisks indicate a significant difference by Rate Ratio test. (E) Microtubule rescue catastrophe rates. (N = 190, 119, 155, and 148 rescues in WT, *spr1, 3x-eb1* and *clasp* mutant backgrounds respectively. Error bars represent SEM. Asterisks indicate significant difference as compared to WT by Rate Ratio test. Significance depicted by * *P* < 0.05, ** *P* < 0.01 and *** *P* < 0.001. Data for all measurements are from 6 cells in 6 plants for each genotype.

## Severing in +TIP mutants

To assess the possible function of +TIPs in modulating severing and in stabilizing microtubule ends after severing, we analyzed the outcomes of microtubule crossover formation in WT and *spr1, 3x-eb1* and *clasp* mutant backgrounds expressing YFP-TUA5 as a microtubule marker. For each crossover, we marked the location, time of creation, time of resolution, whether the crossover was resolved by severing or by depolymerization of either crossover partner, and whether the old microtubule (the preexisting microtubule) or the new microtubule (the microtubule that crosses the preexisting one to form the crossover) at the crossover got severed (See Methods). We also scored if the newly created plus end was shrinking or growing as observed immediately from the crossover site (Fig. 3A and Video 2, Fig. 3B and Video 3, respectively).

**Figure 3.**
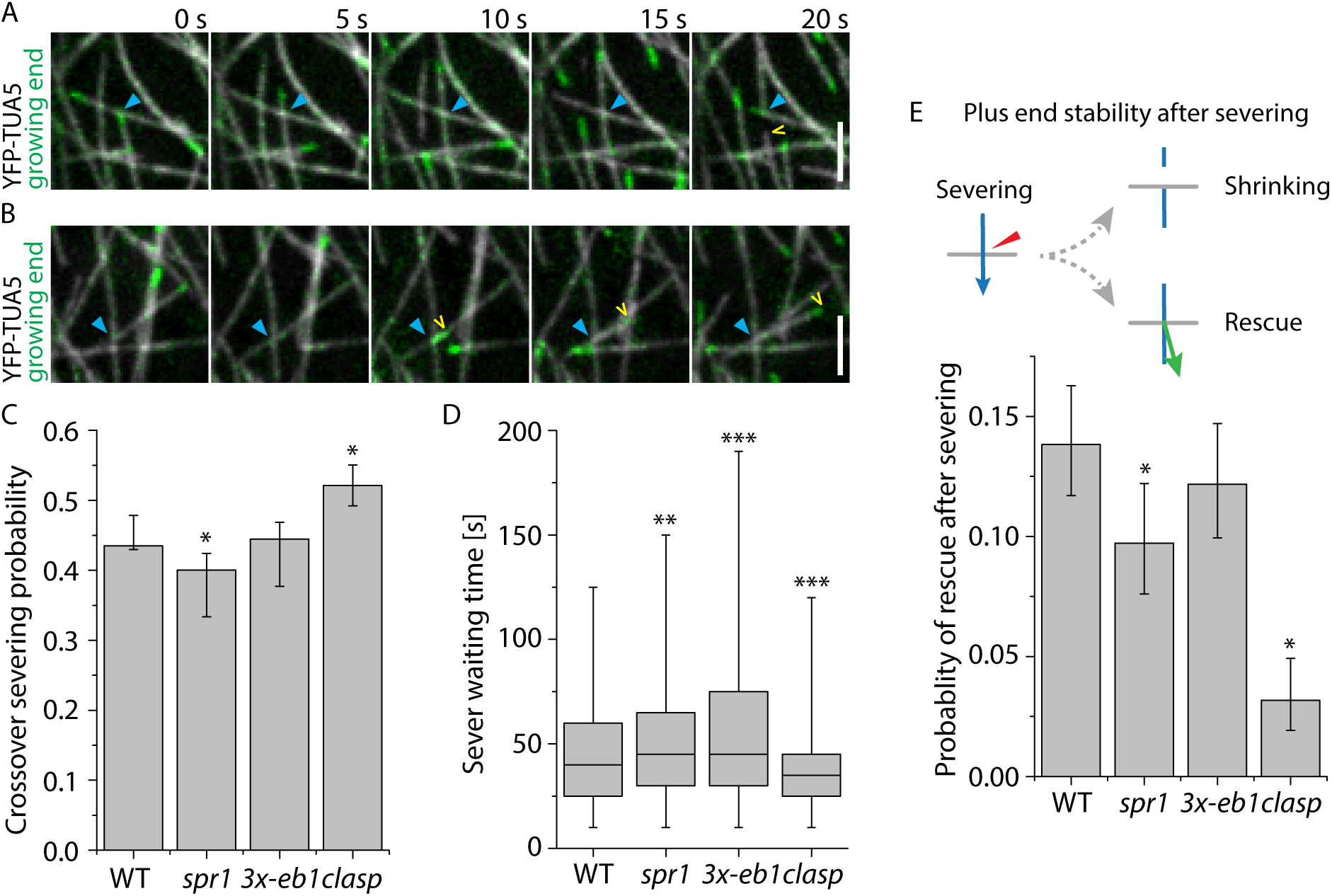
Analysis of microtubule crossovers in +TIP mutants. (A and B) Examples of crossovers (blue arrowhead) where severing occurs, followed by (A) depolymerization (See Video 2) or (B) polymerization (See Video 3) of the new plus end (yellow arrowhead) in a WT cell expressing YFP-TUA5. Scale bars are 3 μm. (C) Severing probability per crossover in cells expressing YPF-TUA5 in WT, *spr1, 3x-eb1*, and *clasp* backgrounds (N = 2027, 1696, 1718, and 1147 crossovers in 6 cells, respectively). Error bars show 95% confidence internals. Asterisks indicate a significant difference from WT by Fisher’s exact test. (D) Waiting times from the observed moment of crossover generation until observed evidence for MT severing in WT, *spr1, 3x-eb1*, and *clasp* backgrounds (N = 2027, 1696, 1718, and 1147 crossovers in 6 cells, respectively). Boxplots show the 25^th^ and 75^th^ percentile as box edges, the line in the box indicates median value and the whiskers show the 2.5^th^ and 97.5^th^ percentile. A Kruskal-Wallis test showed significant differences among reorientation speed distributions among the genotypes (*P* < 0.001). Asterisks indicate significant difference from WT by Mann-Whitney U test. (E) Probability of new plus ends created by severing at crossovers being initially observed in a growing state (N = 882, 679, 764, and 598 crossover severing events in WT, *spr1, 3x-eb1*, and *clasp* backgrounds respectively). Error bars show 95% confidence intervals. Asterisks indicate a significant difference from WT by Fisher’s exact test. Significance depicted by * *P* < 0.05, ** *P* < 0.01 and *** *P* < 0.001.

We observed no significant difference (Fisher’s exact test) in the likelihood of observing severing after crossover formation between WT and *3x-eb1*, with both showing a 44% probability of severing (Fig. 3C). In the *spr1* mutant, we measured a slight decrease in severing probability per crossover as compared to WT, at 42% (*P* < 0.05 Fisher’s exact test). By contrast, the *clasp* mutant showed a 54% probability of severing per crossover, a rate that is markedly higher than WT (*P* < 0.05 Fisher’s exact test) (Fig. 3C).

The severing likelihood depends both on the kinetics of severing itself (e.g. katanin recruitment, activation and action), and on microtubule dynamics, as crossovers get resolved either by severing or by a microtubule end depolymerizing beyond the crossover location. To assess better severing activity itself we measured the time from crossover formation to the time that optical evidence of severing was observed, a quantity we term the sever waiting time (Fig. 3D and Fig. S1). When crossovers are resolved by microtubule dynamics, a sever waiting time is of course not calculated. When comparing the observed sever waiting time distributions we found that both *spr1* (*P* < 0.01 Mann-Whitney U test) and *3x-eb1* (*P* < 0.001 Mann-Whitney U test) have a significantly longer mean sever waiting time compared to WT. Strikingly, the *clasp* mutant shows a significant reduction in sever waiting time for microtubule severing than we found for WT (P < 0.001 Mann-Whitney U test). In summary, microtubule severing activity is increased in the *clasp* mutant, suggesting that CLASP antagonizes severing activity.

## Rescue after severing in +TIP mutants

During blue light-induced microtubule reorientation, most of the new growing microtubules are created by katanin mediated severing (Lindeboom et al., 2013b). An outstanding question is how these new plus ends get stabilized immediately after severing, as they most likely lack the stabilizing GTP-cap. We therefore compared the fates of the new plus end that are created by severing in WT and the +TIP mutants (Fig. 3E). To score severing at crossovers, we asked if there was evidence of either the formation of an optically resolved gap, or a new growing after a new crossover is made. In a previous we study, all such gaps formation and new growing ends emerging from cross overs were found to be dependent on katanin mediated severing (Lindeboom et al., 2013b). Gaps that formed due to microtubule depolymerization were always observed from shrinking of the new plus end. The frequency of stabilization and regrowth of a new plus end was determined by dividing the number of new growing ends emerging from crossovers over by total number of observed severing events (see Methods for further details). In *3x-eb1* mutants, we found that plus ends of microtubules were observed in a growing state 12.2 % of the time immediately following their creation by severing. which was not significantly different from the 13.8 % rate observed in WT (Fisher’s exact test). The probability of plus end regrowth from the crossover site after severing was significantly reduced to 9.7 % (*P* < 0.05 Fisher’s exact test) in the *spr1* mutant, a rate 70% that of WT. Most strikingly, in the *clasp* mutant, we only observed the new plus end created by severing in a growing state 3.2 % of the time (*P* < 0.05 Fisher’s exact test), a rate only 22% that of WT. Thus, these tests of + TIP mutants reveal that SPR1 and CLASP both aid the rescue of plus ends immediately after they are created by severing in epidermal cells during array reorientation, but CLASP is the dominant player, with rescue being severely impaired by loss of CLASP function alone.

## Rescue is associated with CLASP localization but not with crossovers

In higher plants, GFP-tagged CLASP protein localizes to growing plus ends (Kirik et al., 2007) as described in animal cells (Akhmanova et al., 2001), but also accumulates along the lattice of cortical microtubules (Ambrose et al., 2007; Kirik et al., 2007), showing a heterogeneous distribution that includes brighter punctae (Kirik et al., 2007). We confirmed this heterogenous localization pattern along the lattice in the etiolated hypocotyl cells used in this study (Fig. 4A and B, Video 4) and also observed that labeled CLASP accumulates in areas of high microtubule curvature (Fig. 4E, and see especially dynamic examples in Video 5). We speculated that such distributed prepositioning along the lattice might be useful for rescue function when new ends are generated mid-lattice by severing. In support of this possibility, *in vitro* studies of *Schizosaccharomyces pombe* CLASP also showed heterogeneous localization of CLASP along microtubules, and that local CLASP density was correlated with the likelihood of rescue (Al-Bassam et al., 2010). To determine if rescue is influenced by local CLASP concentration *in vivo*, we analyzed the ratio of CLASP to microtubule signal and related this to the behavior of shrinking microtubule ends (Fig. 4C, see Methods). We quantified the CLASP to MT signal ratio in 2716 frames where the microtubule continued shrinking and 301 instances where the microtubule got rescued. When we compared the signal intensity ratio between shrinking and rescue frames we measured a significantly higher CLASP to microtubule ratio at locations where rescue was observed (*P* < 0.001 Mann-Whitney U test, one-tailed) (Fig. 4D), indicating that locations of higher CLASP concentration on the microtubule lattice are preferential rescue sites for interphase microtubules in plants.

**Figure 4.**
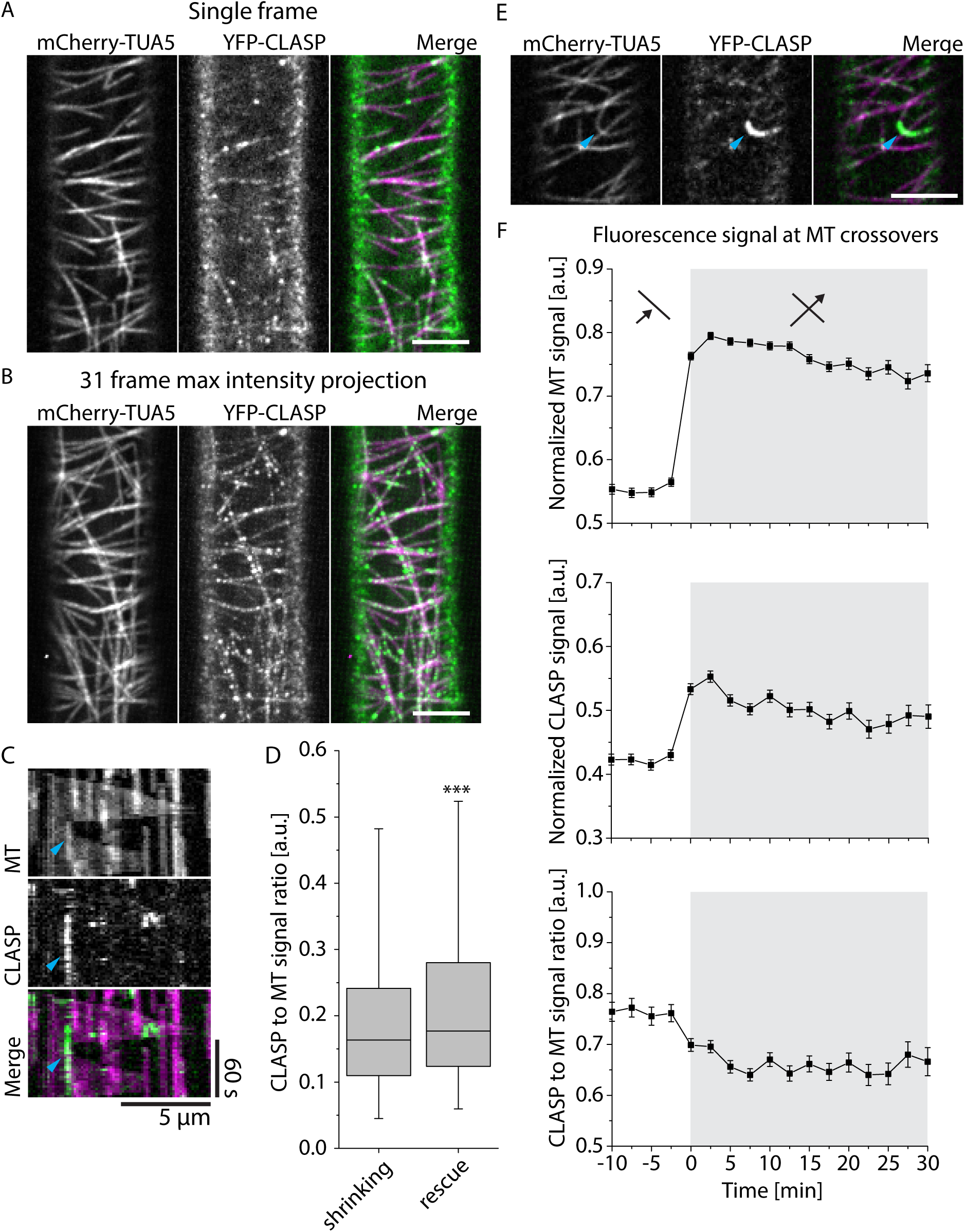
CLASP signal intensities at crossovers and rescue sites. (A and B) Example of MT and CLASP signal distribution in a 3-day old dark grown hypocotyl epidermal cell of a *clasp* mutant that is complemented by a YFP-CLASP construct and in addition expresses mCherry-TUA5 (see Video 4). A single frame is shown in (A) and (B) shows a maximum intensity projection of the first 31 frames at a 5s time interval. Scale bar 5 μm. (C) Kymograph of a MT rescue event coinciding with high CLASP signal intensity. Cyan arrowhead indicates the point of MT rescue. (D) Boxplot of CLASP to MT signal intensity ratio for microtubules that continue shrinking and microtubules that get rescued. Boxplots show the 25 ^th^ and 75 ^th^ percentile as box edges, the line in the box indicates median value and the whiskers show the 2.5^th^ and 97.5^th^ percentile. Microtubules were observed shrinking in 2716 frames and we observed rescue 301 times. CLASP to MT signal intensity ratios were shown to be significantly higher in locations where the rescues occurred by Mann-Whitney U test, significance depicted by * *P* < 0.05, ** *P* < 0.01 and *** *P* < 0.001. (E) Example of high CLASP signal intensity on highly curved microtubule in 3-day old dark grown hypocotyl epidermal cell expressing YFP-CLASP and mCherry-TUA5 (See Video 4). Scale bar 5 μm. (F) MT and CLASP signal intensities during crossover formation. The top panel shows the relative MT signal intensity, the middle panel the relative CLASP signal intensity and the bottom panel shows the ratio of CLASP to MT signal intensity (N = 839 crossovers in 6 cells in 6 plants). The white background indicates the time before the crossovers are formed and the gray background indicates the time at which the crossover is formed.

A recent study reported an increased rescue rate at microtubule crossovers in animal cells (Aumeier et al., 2016). To test if rescues at crossovers are also more likely in *Arabidopsis* we used kymograph analysis to compare the microtubule signal intensity measured at shrinking plus ends to that measured at locations where plus ends were rescued. Microtubule signal intensity is higher at crossovers due to the combined signal of microtubules at crossover sites. If indeed microtubule rescue is elevated at crossovers in plants, we would predict that microtubule signal should be higher on average at sites of rescue as compared to that at locations where plus ends are shrinking. However, we found that the relative microtubule signal intensity was not significantly higher where plus ends were rescued (N = 301) as compared to locations where they continued to shrink (N = 2716) in *Arabidopsis* (*P* > 0.98 Mann-Whitney U test, one-tailed) (Fig. S3). In addition, analysis of the ratio of GFP-CLASP to microtubule signal during crossover formation indicated that CLASP was actually under-represented at crossovers rather than over represented (Fig. 4F, N = 839 crossovers in 6 plants, note standard error bars for CLASP/MT signal before and after crossover). Taken together, these data indicate that CLASP localization on the lattice is correlated with rescue which may help explain the special function of CLASP in plus end rescue follow severing. However, CLASP does not accumulate specifically at crossovers, nor are crossovers themselves hot spots for rescue in plant interphase arrays.

## Simulations for microtubule propagation efficiency by severing in +TIP mutants

Our measurements of microtubule array reorientation revealed that CLASP is a key player in cortical array remodeling in response to light stimulation. Measurements of microtubule behaviors revealed two differences that stood out when comparing *clasp* mutants to WT and the other +TIP mutants: a 28% reduction of the intrinsic rescue rate from 1.56 to 1.12 events/minute, and a 4.7 fold decrease in the probability of plus end growth immediately following severing from 14% to 3%. Significant but quantitatively more modest changes in other parameters of polymer dynamics, such as growth and shrinkage rates, were also observed. To assess the contributions and significance of these changes in polymer behaviors and dynamics in amplifying the new population of microtubules that drives array reorientation, we designed and employed a probabilistic model for generating longitudinal microtubules by severing and rescue (Nakamura et al., 2018). The simulations were parameterized with estimates from our *in vivo* measurements of microtubule dynamics and crossover outcomes in WT, *spr1, 3x-eb1* and *clasp* backgrounds (See Table S1), together with measurements of the spacing between transverse microtubules before reorientation (Fig. S2 and Methods). In each simulation run, a single microtubule was initiated transversely to a grid of pre-existing microtubules and its fate and those of its progeny were assessed as a function of time. For each genotype or experimental test, N = 5 × 10^4^ runs were performed to build up a probabilistic picture of microtubule fate. The results were analyzed in terms of two measures: the probability that a microtubule, and all of its progeny created at severing events, die out (extinction probability), and the expected number of surviving descendants of a microtubule at a fixed time point after initiation (Fig. 5A and B).

**Figure 5.**
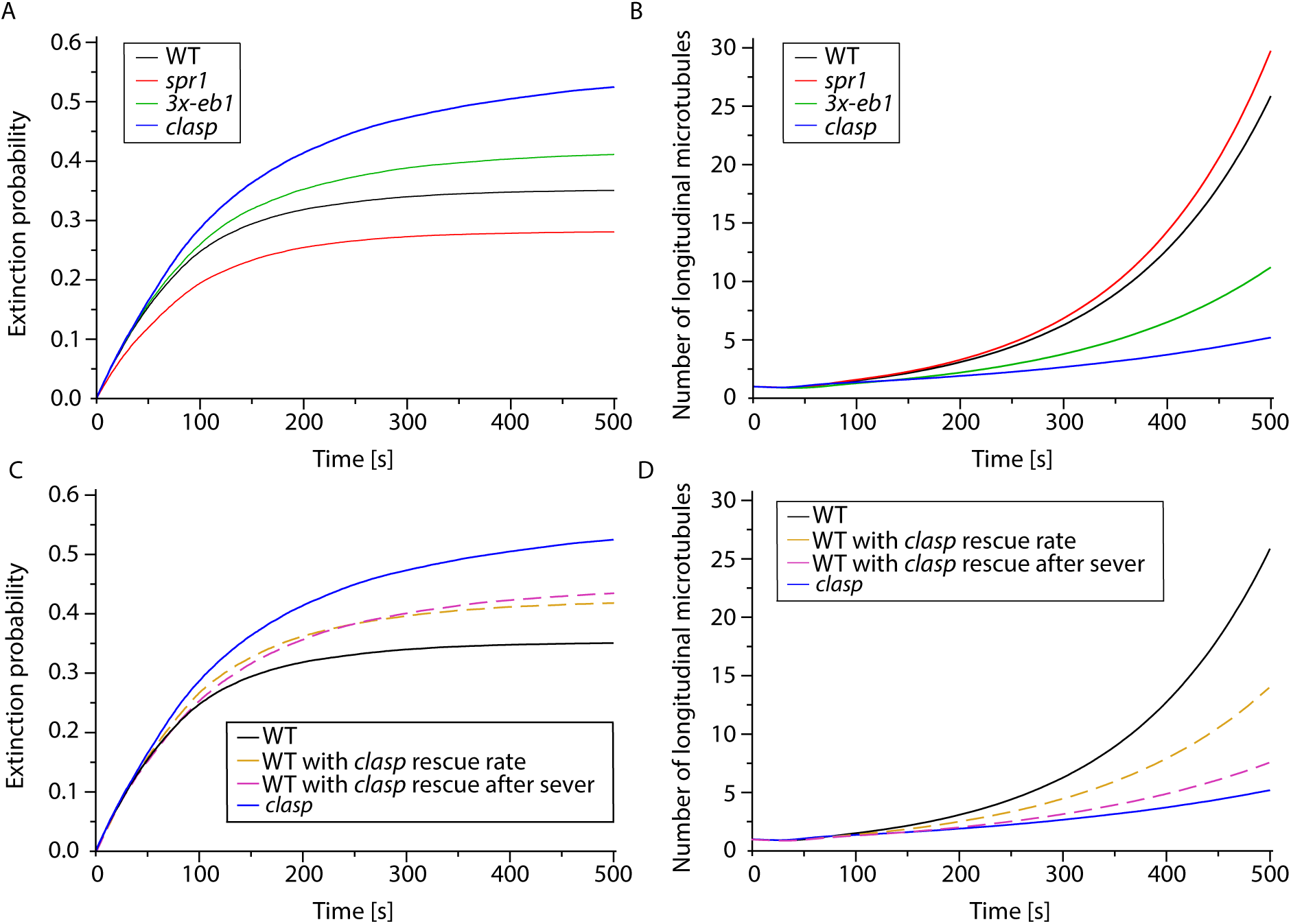
Simulation of microtubule amplification driven by severing. (A) Time evolution of the fraction of extinctions in WT, *spr1, 3x-eb1*, and *clasp* mutants. Extinctions are more likely in less stable mutants, i.e. in mutants that are less deep into the unbounded growth regime. (B) Time evolution of the average number of microtubules during the amplification process shows that a lower probability of rescue after severing and a less deep unbounded growth regime in *3x-eb1* and *clasp* mutants result in a slower amplification as compared to *spr1* and WT. (C) Time evolution of the fraction of extinctions in WT with *clasp* intrinsic rescue rate, and WT with *clasp* rescue after severing probability *in silico* mutants, shows that even if the two reduced rescue parameters have a similar effect on the extinction probability, (D) the probability of rescue after severing *P_s,+_* has a greater impact on the amplification process. Indeed WT microtubules with *clasp P_s,+_* perform an amplification that almost completely reproduces the *clasp* results (dashed purple curve).

For all genotypes, simulations resulted in net microtubule amplification (Fig. 5B) despite a finite probability of failure for each trial (Fig. 5A). This outcome is consistent with experimental observations, where some degree of reorientation was observed for all genotypes, even in the most severe mutant, *clasp*. These results can be understood by examination of the values for the basic microtubules dynamics (growth and shrinkage speeds, transition frequencies), which for all genotypes are in the so-called unbounded growth regime, a state where the measured polymer dynamics predict net polymer growth over time, rather than a state of equilibrium. The measure for this effect is the average growth velocity (Dogterom and Leibler, 1993). A positive average growth velocity itself leads to net amplification because individual microtubules are expected to survive on average in spite of being severed (Tindemans and Mulder, 2010).

While all simulations showed a degree of amplification, the magnitudes of amplification and the probabilities of extinction varied markedly by genotype and mirrored the measured defects on reorientation speed, with *clasp* being the most defective in both amplification and having the highest likelihood of extinction, followed by *3x-eb1* and *spr1* (Fig. 5A and B). Thus, not only do the measured microtubule dynamics and behaviors in *clasp* predict the lowest rate of net amplification, but also the highest likelihood of amplification failure.

In order to understand which of the two major changes in microtubule dynamics observed in the *clasp* mutant might more greatly affect the speed of amplification - the reduction in the intrinsic rescue rate (*r_r_*), or the quantitatively larger reduction in the probability of rescue after severing (*P_s_*_,+_) - we created two synthetic *in silico* mutants: WT with the *clasp* intrinsic rescue rate (*WT/clasp-r_r_*), and WT with the *clasp* probability of rescue immediately after severing (*WT/clasp-P_s,+_*). As shown in Fig. 5C, both mutations have a similarly detrimental effect on extinction probability with respect to WT. However, amplification in *WT/clasp-P_s,+_* is slower than that in *WT/clasp-r_r_*, indicating that the change in *P_s,+_* dominates the amplification defect when CLASP function is lost (Fig. 5D). Further, the reduction in *P_s,+_* by itself almost completely reproduces the *clasp* phenotype. Taken together, the simulation analyses quantify the importance of CLASP function in amplifying new longitudinal microtubules, and support the hypothesis that the dominant function of CLASP in building a new array by severing-driven microtubule amplification is by its specific function in rescue immediately after severing.

## Discussion

In the cortical arrays of higher plants, growing microtubule plus ends are created both by microtubule nucleation (Wasteneys and Williamson, 1989; Nakamura et al., 2010) and by microtubule severing (Wightman and Turner, 2007; Zhang et al., 2013; Lindeboom et al., 2013b). A recent *in vitro* study has shown that templated nucleation, whether from nucleation complexes or microtubule seeds, is an energetically unfavorable process that can be made more efficient by microtubule associated proteins (Wieczorek et al., 2015). The lack of a stabilizing GTP-cap presents a second energetic challenge to the creation of new microtubules from severed ends. Our data identify CLASP as a potent factor for promoting growth from plus ends created by severing ends *in vivo*. This function might therefore be due to lowering the energetic barrier of templated polymer initiation from these ends, by antagonizing the transition to catastrophic disassembly in the absence of a GTP-cap, or perhaps both.

The precise mechanisms by which CLASP might act to stabilize severed ends and promote regrowth have not been identified. Observation of free tubulin at sites of CLASP accumulation *in vitro* suggested that CLASP might act to concentrate tubulin locally to promote rescue of shrinking minus ends (Al-Bassam et al., 2010). An alternative, and not mutually exclusive, possibility is that interaction of CLASP with the microtubule lattice provides stabilizing energy. Proteins that influence microtubule stability often do so based on preferential binding to αβ-tubulin of specific curvature (Brouhard and Rice, 2014). We indeed observed examples of relatively higher CLASP fluorescence along highly curved microtubule regions (Fig. 5b and Video 5). Therefore, an alternative model for CLASP’s action as a rescue factor is that CLASP could halt the progression of curvature that occurs in microtubule protofilaments during depolymerization by binding to tubulin. It is interesting to note that we found that CLASP function antagonizes microtubule severing at microtubule crossovers (Fig. 3C and D). Since microtubule severing requires deformation of the microtubule lattice to break lateral protofilament bonds, it is possible that CLASP offers protection against severing by the same mechanism that stabilizes microtubule plus ends. Finally, the decrease in the likelihood of plus end regrowth following severing in the *clasp* mutant, the function that our modeling studies indicated dominated CLASP function in promoting microtubule amplification by severing, was much greater in relative magnitude than the loss in rescue observed in the general population of rapidly shrinking plus ends. This observation suggests that the mechanism or mechanisms by which CLASP stabilized plus ends and promotes rescue may be more effective at new plus ends just created by severing as compared to plus ends that are already shrinking rapidly.

Severing is a fundamental tool employed by cells across phyla to manipulate and regulate cytoskeletal arrays. Severing is proposed to not only aid in the disassembly of microtubules, but to play an important role in building new arrays in important cell types including neurons, epithelial cells and meiocytes (Roll-Mecak and Vale, 2006). Our recent observations and experiments in higher plant cells provided direct and quantitative evidence that severing can in fact act generatively to build new microtubule arrays, and we report here an important feature of this mechanism - the specific rescue by CLASP of the new plus ends created by severing. In animal cells, the proteins used to stabilize the new plus ends generated by severing have not been identified. As CLASP is conserved across animals and plants, our studies suggest that it may be a compelling candidate to investigate for this function to enable microtubule severing to rapidly build new cytoskeletal arrays.

## Methods

### Plant material

Microtubules were marked by introducing the 35S-YFP-TUA5 construct (Shaw et al., 2003) into four genetic backgrounds of *Arabidopsis thaliana* – Wild type (WT), *spr1–6* (Sedbrook et al., 2004), *cls1–1* (Kirik et al., 2007), *and 3x-eb1 (see below) -* all ecotype Col-0. Wild type (Shaw et al., 2003) and *spr1–6* (Sedbrook et al., 2004) transformants were described previously. For this study 35S-YFP-TUA5 (Shaw et al., 2003) was introduced into *cls1–1* by *Agrobacterium* mediated transformation. In the manuscript, *spr1–6* and *cls1–1* are referred to as *spr1* and *clasp*.

Introduction of 35S-YFP-TUA5 into a triple mutant *eb1* background was accomplished as follows. The *Ateb1a-2* T-DNA insertional mutant (WiscDsLox481–484J12; insertion in the 7th exon of AT3G47690) and the *Ateb1b-2* T-DNA insertional mutant (WiscDsLox331A08; insertion in the 6th intron of At5g62500; see Galva et al., (2014) for additional details) were obtained from the Arabidopsis Knockout Facility (University of Wisconsin Biotechnology Center (Woody et al., 2007)). The *Ateb1c-2* T-DNA insertional mutant (SALK_018475; insertion in the 3rd exon of AT5G67270) was generated at the Salk Institute (Alonso et al., 2003) and obtained from the Arabidopsis Biological Resource Center (ABRC). *Ateb1a-2, Ateb1b-2*, and *Ateb1c-2* mutant plants were cross-pollenated then progeny self-pollenated to generate the *Ateb1a-2 Ateb1b-2 Ateb1c-2* triple mutant. T-DNA insertions were tracked and homozygosity determined by performing PCR analysis (Klimyuk et al., 1993). Primers and primer pairs are listed in Table S2. We refer to this triple eb1 knockout mutant line as *3x-eb1* throughout the manuscript. This *3x-eb1* line was then transformed with the 35S-YFP-TUA5 microtubule marker (Shaw et al., 2003).

For the MT and CLASP dual labelled line, we transformed *clasp1–1* expressing ProAtCLASP-YFP-AtCLASP (Kirik et al., 2007) with a ProUBQ10-mCherry-TUA5 construct. This microtubule reporter was generated by amplifying a 634bp genomic upstream regulatory fragment of UBQ10 from pUBN-GFP-DEST (Grefen et al., 2010). The CaMV 35S promoter was excised from the modified pMDC43 vector encoding mCherry (Kirik et al., 2007) and replaced by the UBQ10 fragment. LR recombinase (Invitrogen) was used to introduce the *Arabidopsis* TUA5 cDNA (Gutierrez et al., 2009) into the modified pMDC43 Gateway binary vector to generate a ProUBQ10-mCherry-TUA5 construct. The binary vector harboring the construct was then introduced into WT Arabidopsis plants (Col-0) by Agrobacterium-mediated transformation using the floral dip method (Clough and Bent, 1998). Transgenic T1 plants were selected for Hygromycin B selection. T2 progeny plants with normal growth and development (with the exception of mild twisting) were used for experiments.

### Plant growth

Seeds were surface sterilized with a 70% ethanol solution, stratified for 3 days at 4 °C, and sown on 1% agar containing Hoagland’s No. 2 salts at pH 5.7. After 1 h of light exposure on a clean bench, plates were wrapped in aluminum foil, set on edge at approximately a 10° angle to vertical, and seedlings were germinated and grown in darkness for 60–72 h at 22 °C.

### Specimen mounting

Foil-covered plates were unwrapped and seedlings were mounted under red safelight conditions to prevent de-etiolation (Paredez et al., 2006). Seedlings were placed gently on a cover slip in sterile water and affixed with a 2-mm thick 1% agarose pad. To reduce specimen drift, the mounted seedling was rested for 20–30 minutes before observation.

### Microscopy

All observations were made in epidermal cells in the upper hypocotyl, a region where cell expansion is rapid in 3 day old etiolated seedlings. The time lapse imaging of the YFP-TUA5 labeled lines was performed at 5s time intervals for a duration of 30 minutes on a CSU-X1 spinning-disk confocal head mounted on a Leica DMI6000B microscope with Adaptive Focus Control (Leica) and a 100x Plan Apo 1.4 N.A. oil immersion objective, as described previously (Lindeboom et al., 2013b). For each image, excitation was supplied by a 488 nm laser for 300 ms at 4.5 mW as measured at the end of optical fiber feeding the spinning disk unit.

We performed imaging of cells dual-labeled with YFP-CLASP and mCherry-TUA5 with a Yokogawa CSU-X1 spinning disk head mounted on a Nikon Eclipse Ti body with a 100X 1.4 N.A. oil immersion objective and perfect focus system as described previously (Gutierrez et al., 2009; Lindeboom et al., 2013a). Exposures were acquired every 2.5s, using a 491nm laser at 8.2 mW to excite YFP-CLASP for 500 ms, and a 591nm laser at 8.2 mW to excite mCherry-TUA5 for 300 ms.

### Reorientation analysis

We used the ImageJ plugin LOCO (Lindeboom et al., 2013b) and a MATLAB script to extract microtubule orientation output to analyze microtubule reorientation after blue light induction. We used a 21-by-3-pixel kernel that we rotate with increments of 9 degrees over a range of 180 degrees. Then for each pixel we assigned the angle for which the kernel contained the highest total fluorescence signal. To limit our measurements to pixels that contained microtubule signal we performed a default threshold in ImageJ and apply this threshold to the angular data. For each angle, we then determined the fraction of the total pixels above the threshold level. From these angular fractions, we also extracted the longitudinal (L) and transverse (T) order parameters based on over- or underrepresentation in the longitudinal and transverse orientation respectively (Lindeboom *et al*, 2013b). To assure that there was sufficient transverse order at the start of stimulation to measure the transition to longitudinal orientation, datasets were selected for analysis where the initial transverse order parameter was lower than 0.1.

### Analysis of microtubule dynamics

The growing and shrinking ends of individual labeled microtubules were visualized by time-phased subtraction of image time series acquired from seedlings expressing YFP-TUA5 as described previously (Lindeboom et al., 2013b). Trajectories of segmented growing ends in selected regions of interest were created blindly using the PlusTipTracker software package (Applegate et al., 2011). These paths were then used to measure microtubule dynamic parameters using a custom pipeline in MATLAB. First, the paths created by PlusTipTracker were used generate kymographs. All kymographs generated were examined, and those showing clear evidence of plus ends growth at least 5 frames were then traced by hand using a custom graphical user interface in MATLAB. Finally, polymerization and depolymerization velocities, as well as the frequency rescue and severing rates, were calculated from the kymograph traces as reported earlier (Nakamura et al., 2018). This protocol allowed for less biased sampling and measurement of end dynamics than what might result from selection by eye.

### Analysis of severing and events at crossovers

To aid in analysis of the formation and resolution of microtubule crossovers in plants expressing YFP-TUA5 in the different genetic backgrounds, we created time-phased and subtracted image series to highlight new microtubule growth, as described previously (Lindeboom et al 2013). Using a custom written environment in MATLAB we created defined regions of interest in each image series in which we exhaustively marked every new crossover, recording the x and y coordinates, the start of crossover formation, the time of crossover resolution, the angles of the old and new microtubules, and whether the old and/or new microtubule at the crossover got severed (Video 6). Severing was tallied either if an optically resolved gap in signal was formed at the crossover, or if new end growth was observed to emerge from the site of the crossover. Our previous analysis showed that all observed gap and new growing end events are created by katanin mediated severing (Lindeboom et al., 2013b). If a severing event took place at the crossover we documented whether the new plus end created by severing was initially shrinking (gap formation) or growing (new growing end as seen as a “comet” in the time-phased images).

This analysis of events at crossovers was also performed on the 2.5 second interval image time series acquired for the analysis of CLASP and tubulin signal, with additional steps to register the two image channels and to correct for photobleaching (Nakamura et al., 2018). In ImageJ (Rasband, 2012), we first registered the images in the images in microtoubule channel using the plugin MultiStackReg. We then used the transformation matrix from this registration to register the CLASP channel. Each channel was subsequently corrected for photobleaching using an exponential fit to total signal intensity of the original images series. We then isolated a 7-by-7-pixel region centered around each of the crossover coordinates. The 1 pixel wide outer border was used for determining the background signal intensity for both CLASP and MT signal. We discarded the highest 12 pixel values of these 28 pixels, as they typically represent the microtubule signal of the two microtubules forming the crossover. We took the median intensity value of the remaining pixels as the local background for the crossover for each individual frame of each crossover. To calculate the crossover signal intensity, we subtracted the background value calculated from the border from the mean intensity of the inner 5-by-5 pixels of the crossover ROI. We then normalized the signal intensity by dividing each individual crossover signal intensity value by the maximum value for each measurement series. The normalized crossover intensities were calculated for both the CLASP and MT signal, and the crossover event intensities were aligned with each other over time based on either the start or end frame of the crossover event. We calculated the CLASP to MT signal ratio by dividing the normalized CLASP intensity by the normalized MT intensity. The standard errors for this ratio estimate were calculated from the error propagation of the standard errors of the separate CLASP and MT signal intensities.

### Quantification of microtubule and CLASP intensity at sites of rescue

For both the microtubule and CLASP channels, we first applied image registration and bleaching correction in ImageJ. For each frame, we normalized the intensity values per frame to the median intensity value after dividing the intensity values into 10 equal sized bins and discarding the values in the highest and lowest bin to remove outliers. We then used these movies to extract kymographs using the method described above in “Analysis of microtubule dynamics”. We traced the edge of the microtubule signal in the kymograph by hand and used a MATLAB script to identify microtubule shrinkage and rescue events. Since CLASP in *Arabidopsis* cells has been reported to be a plus end tracking protein (Kirik et al., 2007), we wanted to exclude any bias of CLASP measurement that may occur due to rescue and subsequent recruitment of CLASP to the growing plus end. We therefore extracted the CLASP and microtubule intensity in a 3 pixel-wide ROI in the two frames immediately before that location is reached by the depolymerizing microtubule end. We assessed the averaged and normalized pixel intensity of these ROIs and sorted them into two groups for comparison: one group in which the shrinking microtubule continues to shrink in the following frame, and the other group where the microtubule is rescued in the following frame.

### Microtubule bundle distances

We measured the distance between transversely oriented microtubules in WT, and in *spr1, 3x-eb1*, and *clasp* mutant backgrounds expressing YFP-TUA5 as described previously (Nakamura *et al*, 2018).

### Stochastic model

To explore the importance of the stabilization of microtubule plus ends generated by severing, we built a stochastic model of longitudinal microtubules undergoing dynamic instability in a grid of stable transverse microtubules (Nakamura et al., 2018). In the previous manuscript, the model addressed protein activities that affected both plus and minus end dynamics. Here, the activities we studied only affected plus ends by experimental measurement, and minus ends were stable by comparison. Thus, in the simulations presented here we kept plus ends dynamic while holding minus ends stable.

Dynamic parameters of the model are the growing speed *v*^+^, the shrinking speed *v*^−^, the catastrophe rate *r_c_* (the rate of transition from the growing to the shrinking state) and the rescue rates *r_r_* (the rate of transition from the shrinking to the growing state). The values of these parameters were measured *in vivo* in hypocotyl epidermal cells undergoing light-driven reorientation (Fig. 2, Table S1). In addition, the model employs the probability of rescue after severing for the plus end, *P_s,+_*. This parameter is the probability that the newly-created plus end grows just after the severing event.

As the sole static model parameter was the distance *d* between neighboring transverse microtubules, this parameter was calculated according to the distribution,

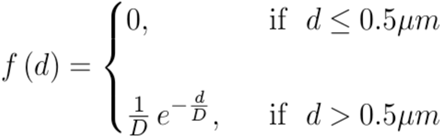
where the length *D* is chosen such that the mean spacing between transverse microtubules *d_avg_* is consistent with the distances experimentally measured between microtubules/bundles (Table S1, Fig. S2).

When the plus end crosses a transverse microtubule, it creates a crossover. At every crossover, there is a competition between two different effects: either the crossover is erased by the shrinkage of the plus end due to dynamic instability, or it leads to a severing event. Whether one effect wins against the other is determined by an intrinsic severing waiting time distribution *W_k_,_θ_(t)*, where *W_k_,_θ_(t)* is the Gamma probability density function (Papoulis, 1984),

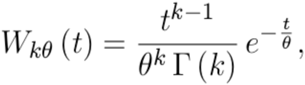
with *k* and *θ* as scale and shape parameters of the distribution, and *Γ*(*k*) as the Euler gamma function (Abramowitz and Stegun, 1965)

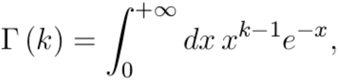

Given the impossibility to measure the intrinsic waiting time distribution for the severing, due to the deletion of some crossovers because of dynamic instability, we have chosen *k* and *θ* such that the distribution of the severing waiting time conditional to the fact that the severing happened, optimally fits the experimentally measured one (see Fig. S4). Given our assumption that the effect of katanin is different for different mutants because of difference in microtubule dynamics and severing properties, the parameters of the distribution are different for WT, *3x-eb1, spr1* and *clasp* mutants. Since we are only interested in the behavior of longitudinal microtubules in the first 500 s of the process, we limit severing to only the longitudinal microtubules.

Every simulation consists of one longitudinal microtubule nucleated in the grid of transverse microtubules, and it lasts for 500 simulated seconds. After that time, the output of the simulation can be either extinction, i.e. the initial microtubule and all its progeny have completely depolymerized due to dynamic instability, or amplification, i.e. there are one or more longitudinal microtubules still alive. We averaged our simulation results over 5 × 10^4^ trials.

The main results of these simulation studies are reported in the results of the manuscript. In order to quantify better the effect of *P_s,+_* and the effect of *r_r_* on the speed of amplification, we performed a sensitivity analysis. In the first case, having WT parameters as background we tuned *P_s,+_*, from 0 to 0.25, and we counted the number of longitudinal microtubules in the simulation after 500s. In the second case we repeated the same procedure, keeping the WT value for *P_s,+_* = 0.15, and tuning the intrinsic rescue rate *r_r_*, from 0.76 events/min to 2.09 events/min. The reason for the choice of these two values for the intrinsic rescue rates is the following: given our interest in the behavior of microtubules in the unbounded growth regime, the rate *r_r_* = 0.76 event/min is the smallest rescue rate required to be in the unbounded growth regime, i.e. it is the value that allows the average growth speed (Dogterom and Leibler, 1993)

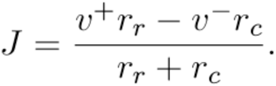
to be non-negative, in particular to be *J* = 0. Taking WT values as our reference values, we can see that [*P_s_*_,+_ = 0.15]: [*P_s_*_,+_ = 0.25] = [*r_r_* = (1.56–0.76) events/min]: [*r_r_* = (2.09–0.76) events/min], motivating our choice to take 2.09 events/min as the intrinsic rescue rate. Fig. S5 shows that the effect on the amplification if we change *P_s,+_* is stronger than if we change *r_r_*, supporting our hypothesis that the key parameter for the speed of amplification is the fraction of rescue after severing.

## Acknowledgements

We thank Zdeněk Lánský for advice on image analysis, David Quint and Luke Rice for helpful discussion. This work was supported by the Carnegie Institution for Science (DWE); NSF grant 1158372 (DWE), and the Human Frontier Science Program (MN). The work of MS is supported by the European Research Council Synergy grant MODELCELL. The work of BMM is part of the research programme of the Netherlands Organisation for Scientific Research (NWO).

## Author contributions

DWE, JJL, MN, VK, TK and AMCE designed experimental strategy. JJL, MN and AH carried out experiments. JJL, MN and DWE analyzed data. BM and MS designed and performed the simulations. JJL, DWE, MN, BM and MS wrote the manuscript. AW and JS provided biological materials. All authors contributed to editing the manuscript.

---

**Figure S1.**
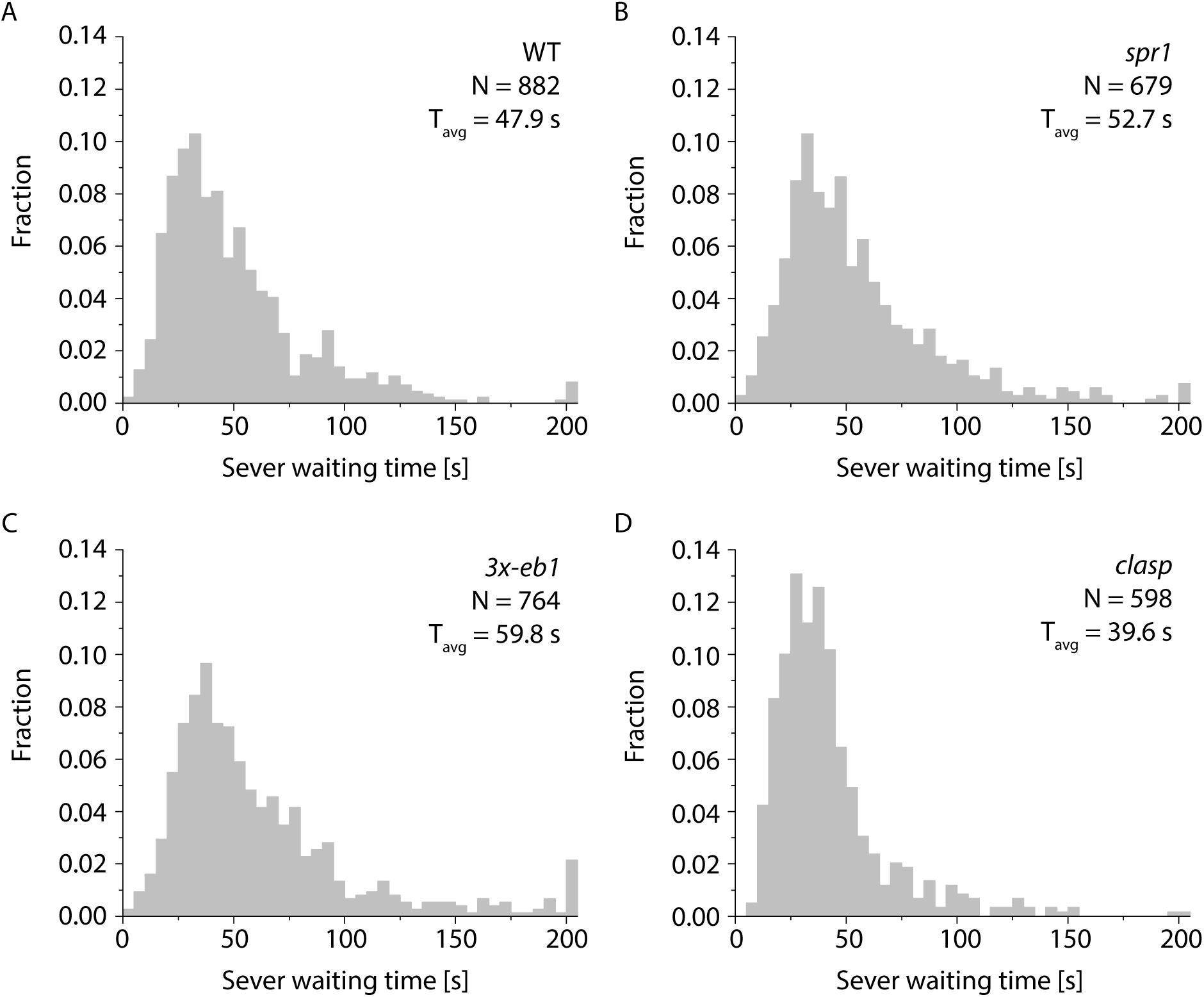
Sever waiting time distributions and severing preferences. (A-D) Distribution of sever waiting times from the moment the crossover is formed in (A) WT, (B) *spr1*, (C) *3x-eb1* and (D) *clasp*. Six plants were used for each genotype. Sever waiting times > 200 s are added up and shown in the last bin.

**Figure S2.**
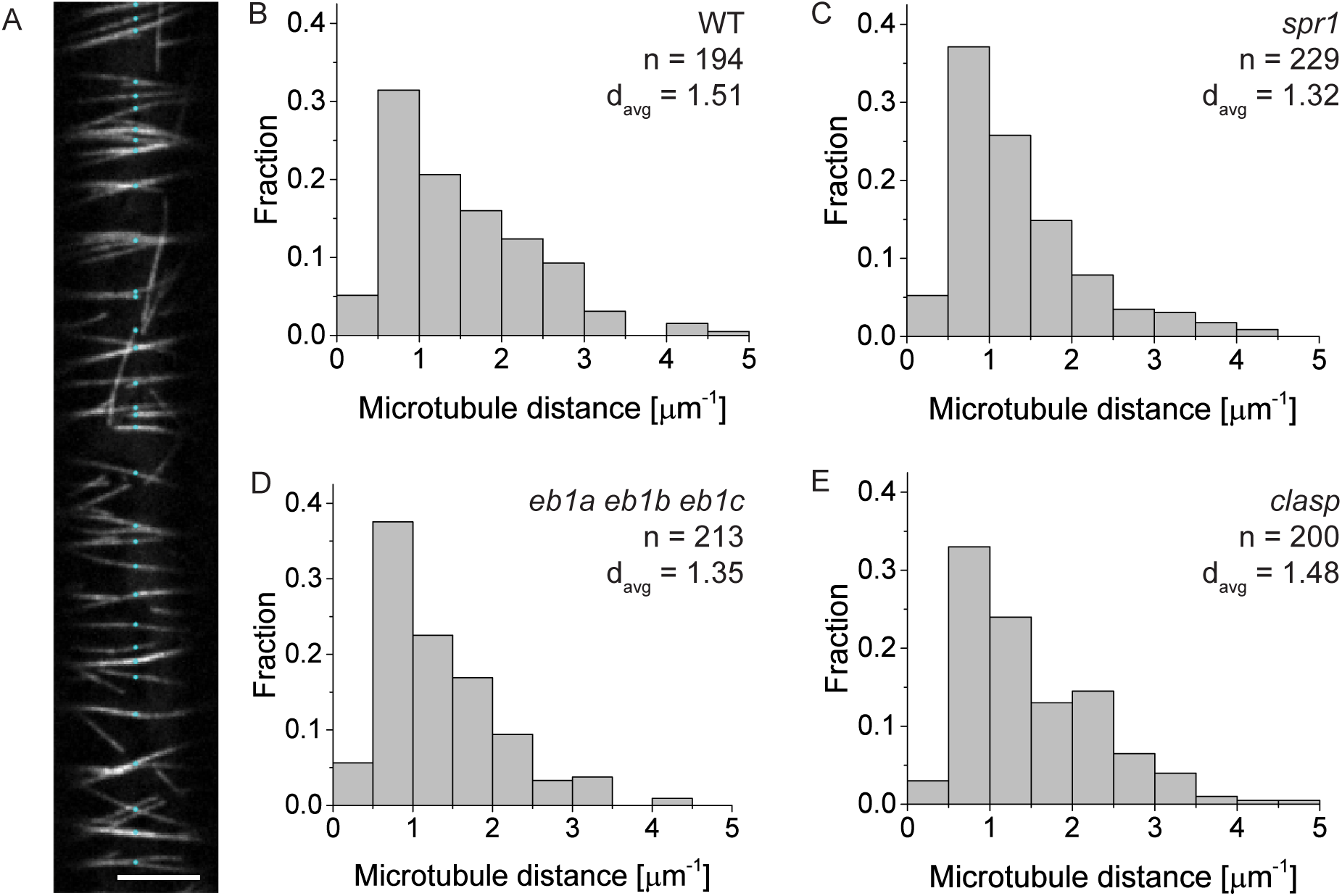
Longitudinal microtubule distances. (A) WT example image with detected microtubules labeled in cyan. Scale bar 5 μm. (B-E) Histogram of microtubule distances in (B) WT, (C) *spr1*, (D) *3x-eb1* and (E) *clasp*. Number of detected microtubules indicated by N, d_avg_ is the average distance between microtubules in μm, we used 6 cells per genotype. A Kruskal-Wallis test followed by a Mann-Whitney U test shows that the distribution of bundle distances only differs significantly from WT for *spr1* (*P* < 0.05).

**Figure S3.**
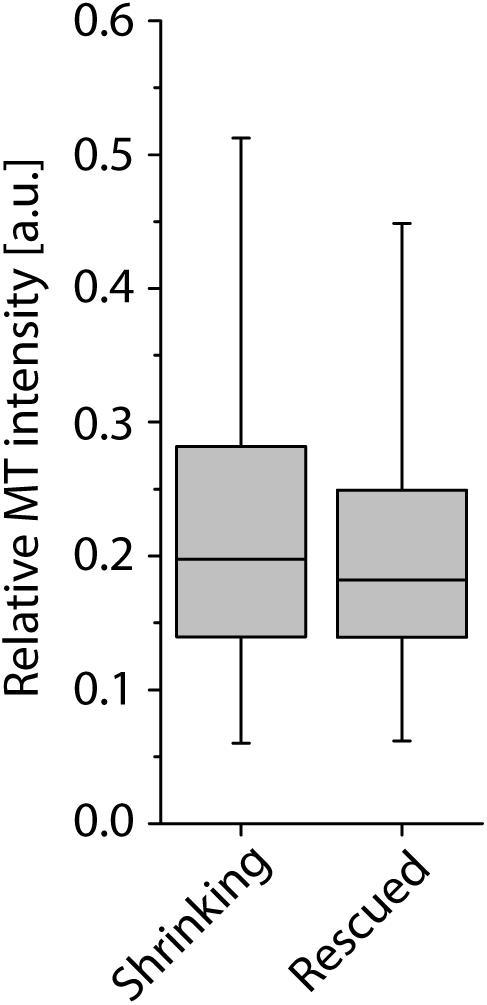
Microtubule signal does not positively correlate with rescue events. Boxplot of relative MT signal intensity on microtubules that continue shrinking and microtubules that get rescued. Boxplots show the 25^th^ and 75^th^ percentile as box edges, the line in the box indicates median value and the whiskers show the 2.5^th^ and 97.5^th^ percentile. Microtubules were observed shrinking in 2716 frames and we observed rescue 301 times. The relative MT signal was not significantly higher for instances of microtubule rescue (p > 0.98 Mann-Whitney U test).

**Figure S4.**
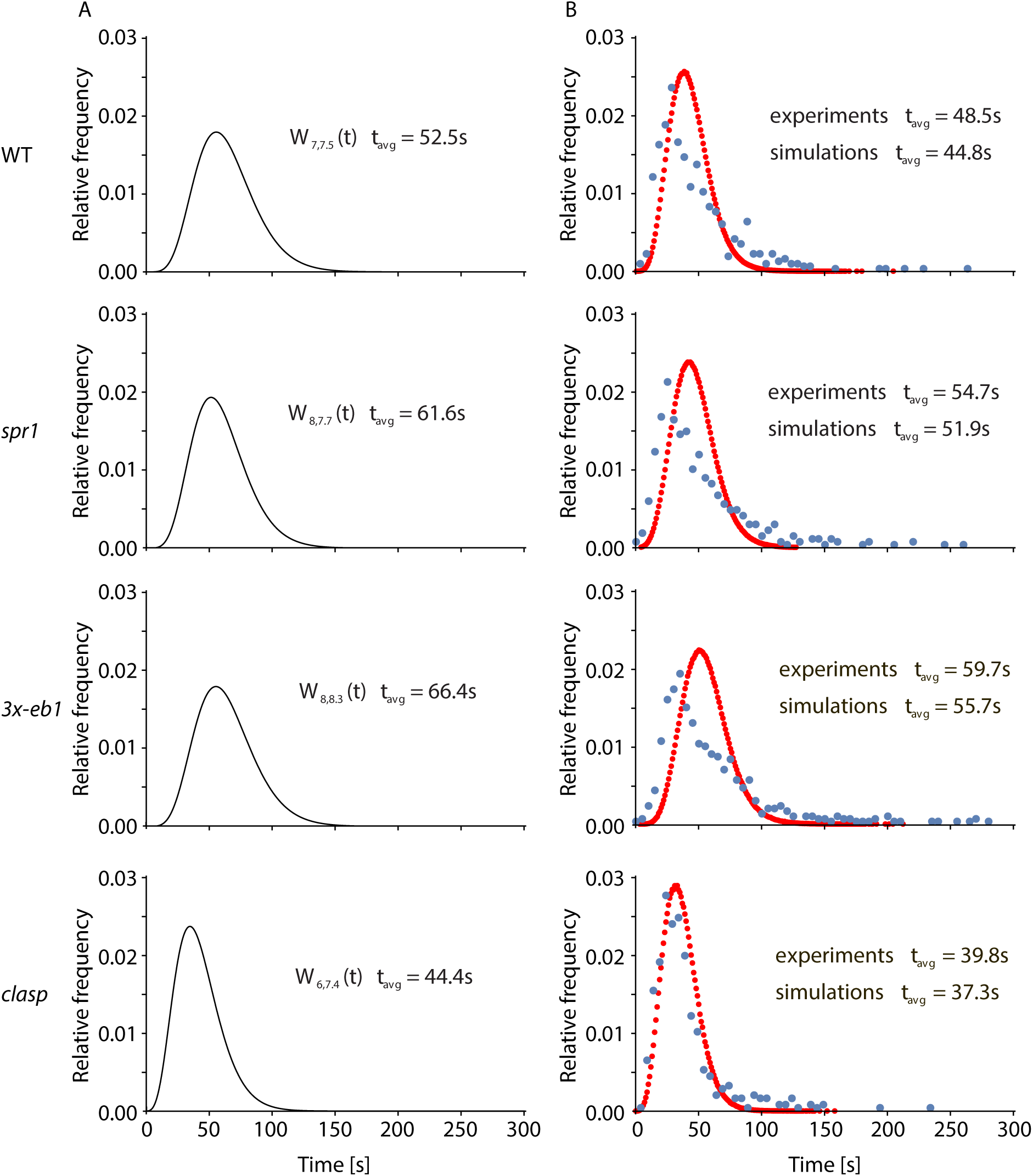
The optimal intrinsic severing waiting time distribution used in simulations (A), and the comparison between computed conditional severing waiting time distribution (red dots) and experimentally measured distribution (blue dots), for WT, *spr1, 3x-eb1*, and *clasp* (B).

**Figure S5.**
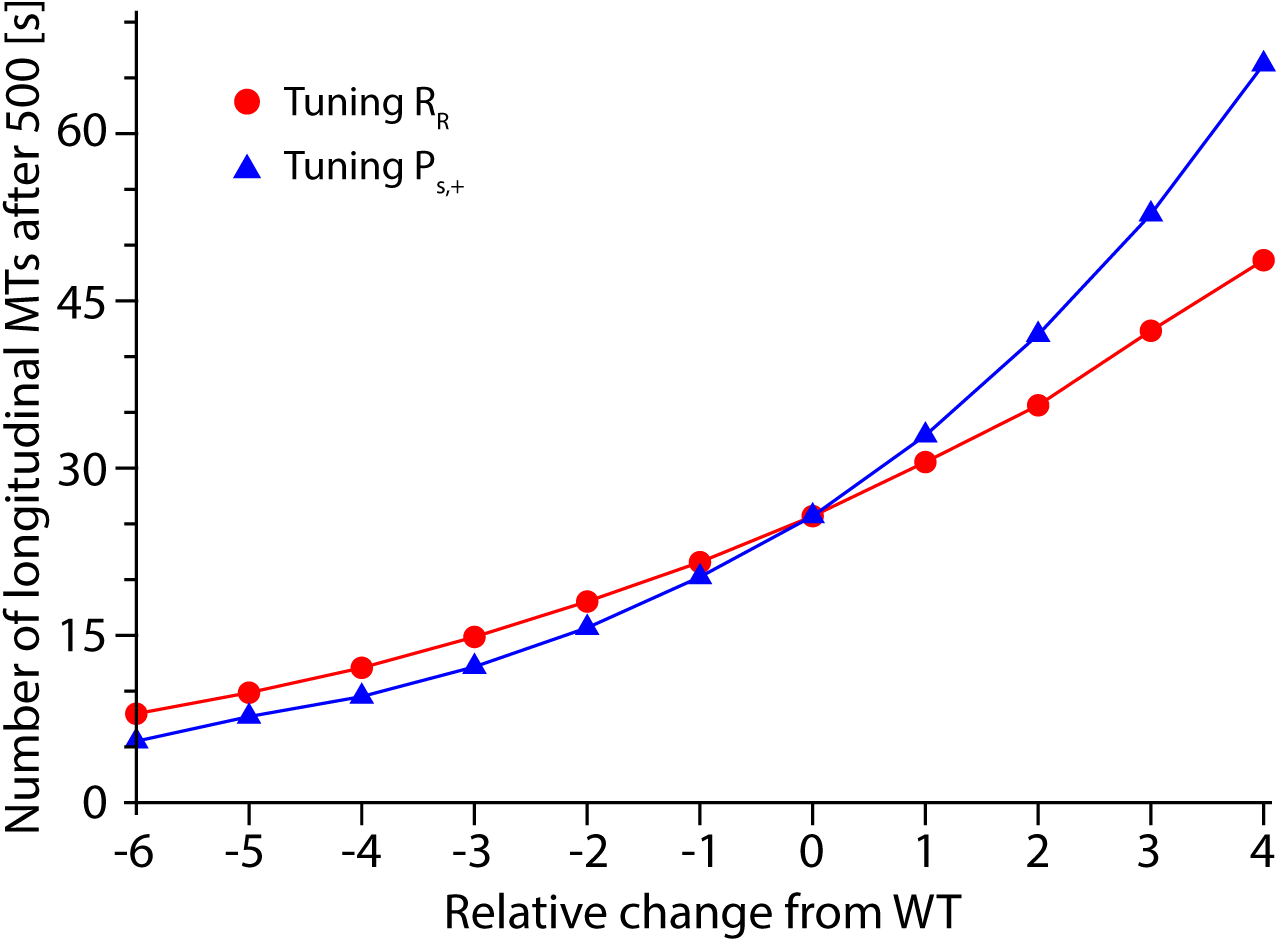
Number of longitudinal microtubules in the simulations after 500s as *P_s,+_* and *r_r_* are varied. Blue triangles represent WT microtubules with changed rescue after severing probability, respectively *P_s,+_* = 0, *P_s,+_* = 0.025, *P_s,+_* = 0.05, *P_s,+_* = 0.075, *P_s,+_* = 0.1, *P_s,+_* = 0.125, *P_s,+_* = 0.15, *P_s,+_* = 0.175, *P_s,+_* = 0.2, *P_s,+_* = 0.225, and *P_s,+_* = 0.25. Red dots represent WT microtubules with changed intrinsic rescue rate, respectively *r_r_* = 0.76 events/min, *r_r_* = 0.89 events/min, *r_r_* = 1.03 events/min, *r_r_* = 1.16 events/min, *r_r_* = 1.30 events/min, *r_r_* = 1.43 events/min, *r_r_* = 1.56 events/min, *r_r_* = 1.69 events/min, *r_r_* = 1.83 events/min, *r_r_* = 1.96 events/min, and *r_r_* = 2.09 events/min.

**Table S1:** Simulation parameters

| Parameter | Description | WT | *spr1* | *3x-eb1* | *clasp* | Units |
| --- | --- | --- | --- | --- | --- | --- |
| $v^+$ | plus end growth velocity | 6.2 | 6.6 | 6.1 | 7.0 | $\mu\text{m min}^{-1}$ |
| $v^-$ | plus end shrinking velocity | 13.5 | 12.4 | 14.9 | 13.5 | $\mu\text{m min}^{-1}$ |
| $r_r$ | rescue rate | 1.56 | 1.57 | 1.42 | 1.12 | $\text{min}^{-1}$ |
| $r_c$ | catastrophe rate | 0.35 | 0.29 | 0.34 | 0.37 | $\text{min}^{-1}$ |
| $d_{avg}$ | average microtubule spacing distance | 1.51 | 1.32 | 1.35 | 1.48 | $\mu\text{m}$ |
| $P_{s,+}$ | probability of rescue after severing | 0.15 | 0.10 | 0.13 | 0.03 | - |
| $k$ | shape parameters of gamma distribution | 7 | 8 | 8 | 6 | - |
| $\theta$ | scale parameters of gamma distribution | 7.5 | 7.7 | 8.3 | 7.4 | s |
| $D$ | mean microtubule spacing | 1.01 | 0.82 | 0.85 | 0.98 | $\mu\text{m}$ |

**Supplemental Table S2.**
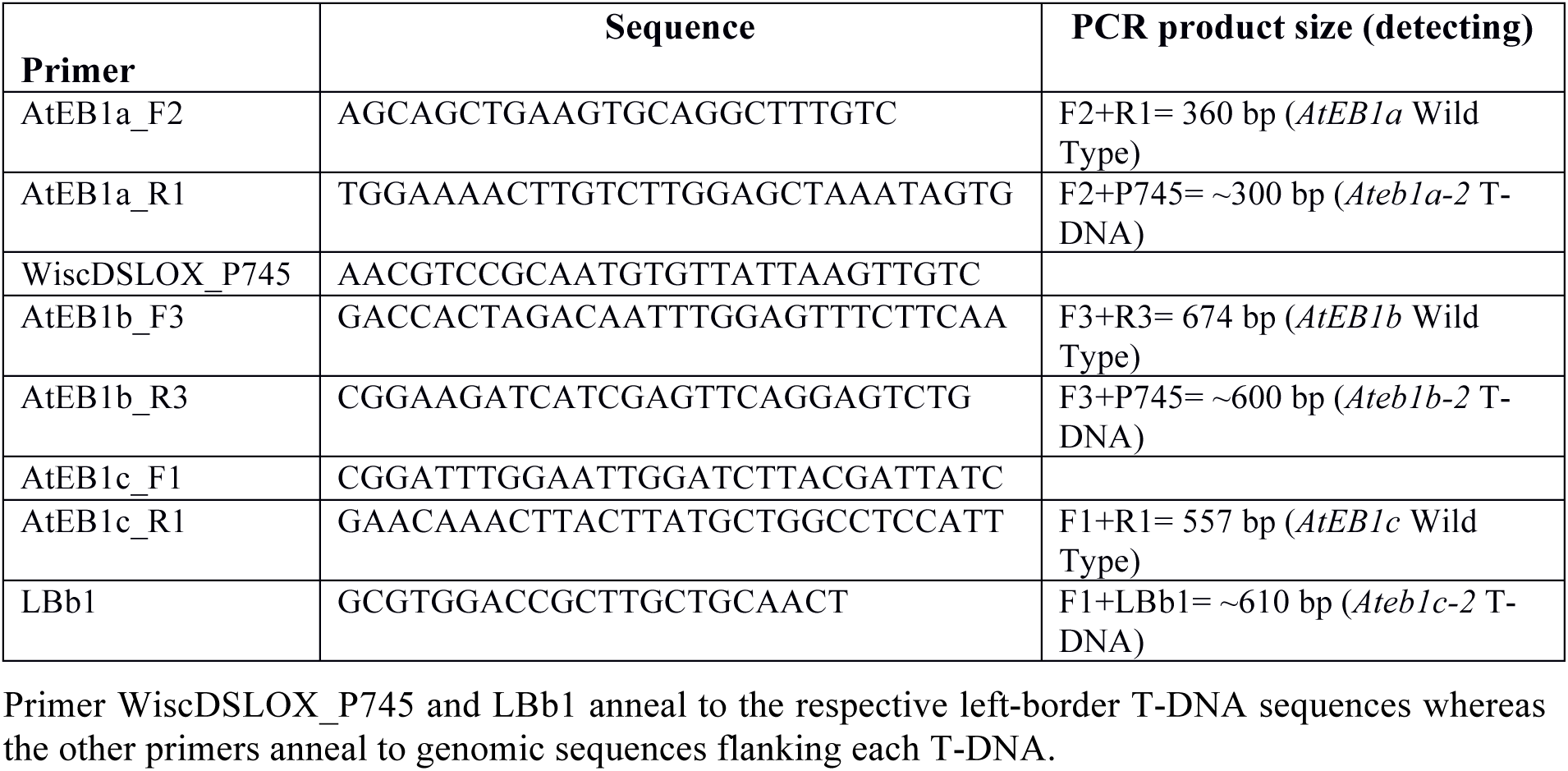
Primers used to score by PCR the presence or absence of corresponding T-DNA insertions in *Ateb1a-2* (WiscDsLox481–484J12), *Ateb1b-2* (WiscDsLox331A08), and *Ateb1c-2* (SALK_018475) segregants.

## Supplementary movie legends

Video 1: Blue light induced microtubule reorientation in dark grown hypocotyl epidermal cells expressing YFP-TUA5 in WT, *spr1, 3x-eb1* and *clasp* genetic backgrounds.

Video 2: Example of crossover severing event in a WT dark grown hypocotyl epidermal cell that generates a shrinking new plus end. The cyan arrowhead marks the microtubule crossover site. The yellow ‘>’ marks the new plus end. Growing microtubule ends are shown in green.

Video 3: Example of crossover severing event in a WT dark grown hypocotyl epidermal cell that generates a growing new plus end. The cyan arrowhead marks the microtubule crossover site. The yellow ‘>’ marks the new plus end. Growing microtubule ends are shown in green.

Video 4: Time lapse movie of CLASP and MT colocalization in dark grown hypocotyl epidermal cell in a *clasp* mutant rescued with a YFP-CLASP construct and co-expressing mCherry-TUA5.

Video 5: Example time lapse Video showing strong CLASP label on highly curved microtubules. Highly curved microtubules are marked by a cyan arrowhead. The example movie features a dark grown epidermal hypocotyl cell of a *clasp* mutant rescued with a YFP-CLASP construct and co-expressing mCherry-TUA5.

Video 6: Example of a section of a WT cell expressing YFP-TUA5 dark grown hypocotyl cell where all the identified crossovers are marked with a yellow ‘+’-sign. Growing microtubule ends are shown in green.

